# Whole-Tissue Distribution Analysis for Visualization of Nanoplastics in the Mouse Brain

**DOI:** 10.64898/2025.12.23.696138

**Authors:** Yang Mi, Tomohiro Ito, Kosuke Tanaka, Osamu Udagawa, Masaki Kakeyama, Yasuo Tsutsumi, Fumihiko Maekawa

## Abstract

Nanoplastics (NPs) are increasingly recognized as environmental contaminants and have been detected in diverse biological tissues; however, their biodistribution in complex organs remains poorly understood. Conventional section-based imaging restricts spatial context and volumetric analysis, making it challenging to map NP distribution in structurally complex organs such as the brain, even with fluorescent labeling. To address these limitations, we developed an integrated workflow combining tissue optical clearing (SeeDB2G) with light-sheet fluorescence microscopy (LSFM), enabling three-dimensional visualization of fluorescently labeled polystyrene (PS) NPs in neonatal mouse brains. At postnatal day 0 - a critical window of heightened vulnerability due to immature barrier systems and rapid neurodevelopment - pups were orally administered spherical PSNPs of 50 or 500 nm. Fluorescence stereoscopic imaging revealed pronounced organ-level accumulation of 50 nm PSNPs in the intestine, liver, kidney, and brain, compared with markedly lower signals from 500 nm PSNPs 24 h post-exposure. Optical clearing rendered the entire brain transparent while preserving fluorescence, allowing LSFM to accurately assess regional PSNP accumulation without sectioning. This workflow enabled whole-brain visualization of size-dependent NP uptake, with 50 nm PSNP detected throughout the brain and the highest accumulation observed in the thalamus and brainstem. Although the current implementation relies on fluorescent labeling and model NPs, this approach provides a scalable platform for whole-organ biodistribution analysis and lays the foundation for mechanistic studies of barrier permeability, developmental vulnerability, and organ-specific interactions.

## 1. Introduction

Nanoplastics (NPs; <1 µm) (Gigault et al., 2018) originating from environmental degradation of larger plastic debris or direct release of engineered polymers, have emerged as pervasive contaminants across aquatic (Uddin et al., 2022), terrestrial (Ullah et al., 2022), and atmospheric systems (Allen et al., 2022). These particles can penetrate biological barriers and accumulate in internal tissues after ingestion, inhalation, or dermal contact. Of these, oral ingestion (Tsochatzis et al., 2024) is considered the predominant route because of the higher estimated intake from contaminated food and water, and experimental oral NP administration in mice demonstrated systemic absorption.

In aquatic invertebrates and fish, polystyrene (PS) NPs accumulate in the intestines (Habumugisha et al., 2023), impairing growth and survival (Feng et al., 2022). In mammals, PS, polyethylene (PE), and polypropylene (PP) fragments have been detected in the liver, kidney, and brain (Lee et al., 2025), often localized near vascular walls (Leslie et al., 2022). A recent autopsy study reported NP accumulation in decedent human brains (Nihart et al., 2025), with markedly higher levels in individuals diagnosed with dementia, raising concerns about potential links to neurological disorders (Eisen et al., 2024) and underscoring the broader relevance of this issue to public health. Beyond the brain, recent studies have confirmed the presence of micro- and nanoplastics (MNPs) in the placenta (Garcia et al., 2023; Halfar et al., 2023), amniotic fluid (Halfar et al., 2023), meconium (Zhu et al., 2024), infant feces, breast milk (Ragusa et al.,2022; Saraluck et al., 2024), and infant formula (Liu et al, 2023), highlighting pregnancy and infancy as sensitive exposure windows. Notably, higher MNP concentrations in preterm births correlate with gestational age and birth weight (Jochum et al., 2025), suggesting potential links with adverse developmental outcomes. Postnatally, breast milk represents a major exposure route, and microplastics (MPs) (Ragusa et al.,2022) and other nanoparticles (Yang et al., 2025) have been detected in human milk. These findings underscore the need to examine early life exposure pathways and their implications in neurodevelopment.

Fluorescently labeled NPs are widely used to study the potential impact of NPs, especially in complex regions such as the brain, as they enable nanoscale particle detection using optical imaging techniques. Administering fluorescently labeled NPs to rodent models has revealed diverse neurobiological consequences of NP exposure, particularly via 20 (Wen et al., 2024)–50 nm PS, with evidence of blood– brain barrier (BBB) penetration, microglial activation (Shan et al., 2022), and behavioral deficits (Paing et al.,2024). Chronic NP ingestion also disrupts the gut microbiota and intestinal barrier integrity (Zhang et al., 2023), suggesting systemic immune involvement, even when brain endpoints are not directly assessed. Emerging evidence also highlights prenatal vulnerabilities. Maternal exposure to PSNPs induces fetal growth restriction (Chen et al., 2022), disrupts postnatal behavioral milestones, and alters brain structure (Harvey et al.,2023), as well as metabolic brain features in offspring (Mercer et al., 2023). Importantly, combined prenatal and postnatal exposure studies have revealed that maternally ingested PSNPs can reach the brain of offspring through postnatal transfer via breast milk, resulting in detectable brain accumulation and neurodevelopmental alterations (Jeong et al.,2021). This finding parallels the evidence of MPs in human breast milk and underscores postnatal transfer as a critical exposure route. Collectively, these results highlight that the neonatal period—characterized by ongoing neurogenesis and immature barrier systems— is a key window for NP uptake and potential neurotoxicity, reinforcing the need for whole-brain mapping approaches to understand biodistribution during this stage.

Despite growing concerns about NP toxicity and increasing reports on their presence across various organs, mapping their distribution in structurally complex regions remains technically challenging, particularly when whole-organ resolution is required to capture spatial dynamics, as in brain-focused studies. Light-sheet fluorescence microscopy (LSFM) offers rapid volumetric imaging with minimal photobleaching, making it ideal for mapping nanoparticle distribution in organs; however, optimal performance depends on the optical transparency achieved via tissue-clearing techniques. Among the available methods (such as CLARITY, CUBIC, and ScaleS), SeeDB2G is an immersion-based and index-optimized protocol that is nonhazardous and faster than many alternatives. It maintains sample size without deformation and minimizes tissue fragility without compromising nanoscale architecture and fluorescence signals. By reducing both light scattering and spherical aberrations, SeeDB2G enables higher-resolution imaging and reliable detection of nanoparticles with fine structural details (Ke et al., 2016).

By integrating fluorescently labeled NPs, the tissue optical clearing technique SeeDB2G, and LSFM, we aimed to establish a workflow that enabled three-dimensional (3D) visualization of NP distribution in neonatal mouse brains. This study focused on the early postnatal period, a key window of neurodevelopment that remains underexplored in NP research. Our approach preserves anatomical integrity and fluorescence signals, allows the mapping of NP accumulation, and offers a methodological framework for future studies on NP biodistribution and developmental vulnerability.

## 2. Materials and methods

### 2.1 NPs used in this study

Carboxylated PS NPs (PSNP) (Polysciences, USA) of two distinct sizes (50 and 500 nm), labeled with a yellow-green fluorescent dye (excitation, 441 nm; emission, 486 nm), were used to assess bioaccumulation. Based on the experimental design, the PSNP stock solution was diluted to a final concentration of 2.5 or 12.5 mg/mL using distilled water (DW) (NIPPON GENE, Japan). The surface morphology of the PS particles was analyzed via scanning electron microscopy (SEM; JSM-7800F, JEOL Ltd., Tokyo, Japan) at magnifications ranging from 10,000× to 50,000× and an accelerating voltage of 5.0 kV, as previously described (Tanaka et al., 2023).

### 2.2 Animals and neonatal treatment

The animal experiment protocol (No. AE-25-07) used in this study was approved by the Animal Use and Care Committee of the National Institute of Environmental Studies (NIES), Japan. The experiments were conducted in strict accordance with NIES guidelines. Pregnant C57BL/6J mice (embryonic day 13–14, E13-14) were purchased from SLC Japan and housed under standard laboratory conditions with unrestricted food and water access. On postnatal day 0 (P0), neonatal mice were weighed and randomly assigned to receive a 30 μL oral gavage of either DW or PSNP at varying concentrations, using a polypropylene feeding tube (FTP-22-25, INSTECH, USA). To facilitate identification, the paw pads of each pup were labeled using a tattooing system (AIMS, USA). The pups were returned to the dam and left undisturbed until sample collection.

### 2.3 Sample collection and whole mouse brain clearing

At 24 h post-treatment, neonatal mice underwent hypothermia-induced anesthesia via ice placement. Perfusion was performed using 8 mL of phosphate-buffered saline (PBS), followed by 8 mL 4% paraformaldehyde (PFA). Tissues of interest were then dissected and immersed in 4% PFA overnight at 4°C for optimal fixation. Subsequently, the samples were either processed for tissue clearing or transferred to PBS containing 0.02% sodium azide for long-term storage.

Neonatal brain clearing was conducted using the SeeDB2 Trial Kit (FUJIFILM Wako, Japan) following the manufacturer’s protocol, with the following optimizations to enhance transparency and preserve anatomical integrity. After washing fixed brain samples in PBS (3 × 5 min), they were transferred to a permeabilization solution (3 mL per sample in 5 mL tubes) and incubated for 18 h. Sequential clearing was performed by incubating samples in Clearing Solution 1, 2, and 3 for 7 h, 17 h, and 24 h, respectively. All the incubation steps were performed at 25°C with gentle shaking. Throughout the process, the brains were handled using ring-tip forceps to minimize tissue deformation. The cleared tissues were mounted in SeeDB2G solution for LSFM. For long-term storage, 0.01% sodium azide was added to the SeeDB2G solution to prevent microbial growth.

### 2.4 Imaging and analysis

Whole-tissue images were acquired using a Leica S8 APO-LED2500 stereo microscope (Leica, Germany) for bright-field imaging and with the addition of a Royal Blue fluorescence adapter equipped with a 500 nm long pass emission filter (NIGHTSEA, USA) for fluorescence imaging.

Light-sheet imaging: Whole-brain 3D imaging was acquired using a Zeiss Light-sheet 7 microscope (ZEISS, Germany) with an 8.36 μm light-sheet thickness in single-illumination mode. The samples were mounted on the holder and immersed in glycerol-based SeeDB2G (RI = 1.46) to ensure optimal refractive index matching. Imaging was conducted using a 488 nm excitation laser with a 5× illumination lens (0.1 NA, ZEISS, Germany) and a 2.5× detection objective (Fluar 0.12 NA, ZEISS, Germany). All image acquisition, processing, and analyses were performed using Arivis Vision4D (ZEISS, Germany). The samples were visualized, stitched, and rendered into 3D reconstructions within the Vision4D workflow environment, with segmentation and volumetric quantification performed using built-in analysis tools.

Quantitative analysis was conducted by selecting three spherical regions of interest (ROIs; 100 µm diameter) within each brain region - cerebellum, cortex, hindbrain, and thalamus - for every sample. To ensure consistency and minimize the impact of hemispheric signal bias - likely due to single-sided illumination in the light-sheet imaging setup - all ROIs showing stronger fluorescence signals across samples were selected from the hemisphere. For each brain region, the mean fluorescence intensity was calculated from the three ROIs, and these values were used for statistical comparisons between the experimental groups.

### 2.5. Statistical analysis

All data are expressed as means ± standard deviations. Statistical analyses were performed using the GraphPad Prism ver. 10.6.1 (GRAPHPAD Software, USA). Group differences were assessed using one-way or two-way analysis of variance (ANOVA) with Tukey’s post-hoc test. Statistical significance was set at p < 0.05.

## 3. Results

### 3.1 Size-dependent biodistribution in neonatal mice

To examine NP uptake and biodistribution in neonatal mice, and to evaluate the impact of particle size, we administered 30 μL 12.5 mg/mL PSNP with two distinct diameters—50 and 500 nm—to P0 mice via oral gavage (Figure 1A). SEM analysis confirmed uniform spherical morphology for both particle sizes, as previously shown (Ito et al., 2025). Although the litter sizes were sometimes insufficient to cover all groups and sexes, we assigned pups such that most dams contributed to the control group and at least one experimental group, including both sexes. Overall, the study included 18 (11 dams), 25 (10 dams), and 16 (7 dams) pups in the DW, 50 nm PSNP, and 500 nm PSNP groups, respectively. Twenty-four hours post-administration, pup survival rates were 68% in the 50 nm PSNP group and 87% in the 500 nm PSNP group, compared to 100% in the control group (Figure 1B). As survival outcomes reflect a combination of acute mortality and maternal behaviors, such as cannibalism and infanticide, which were frequently observed, they are insufficient for generating reliable lethality statistics.

**Figure 1.**
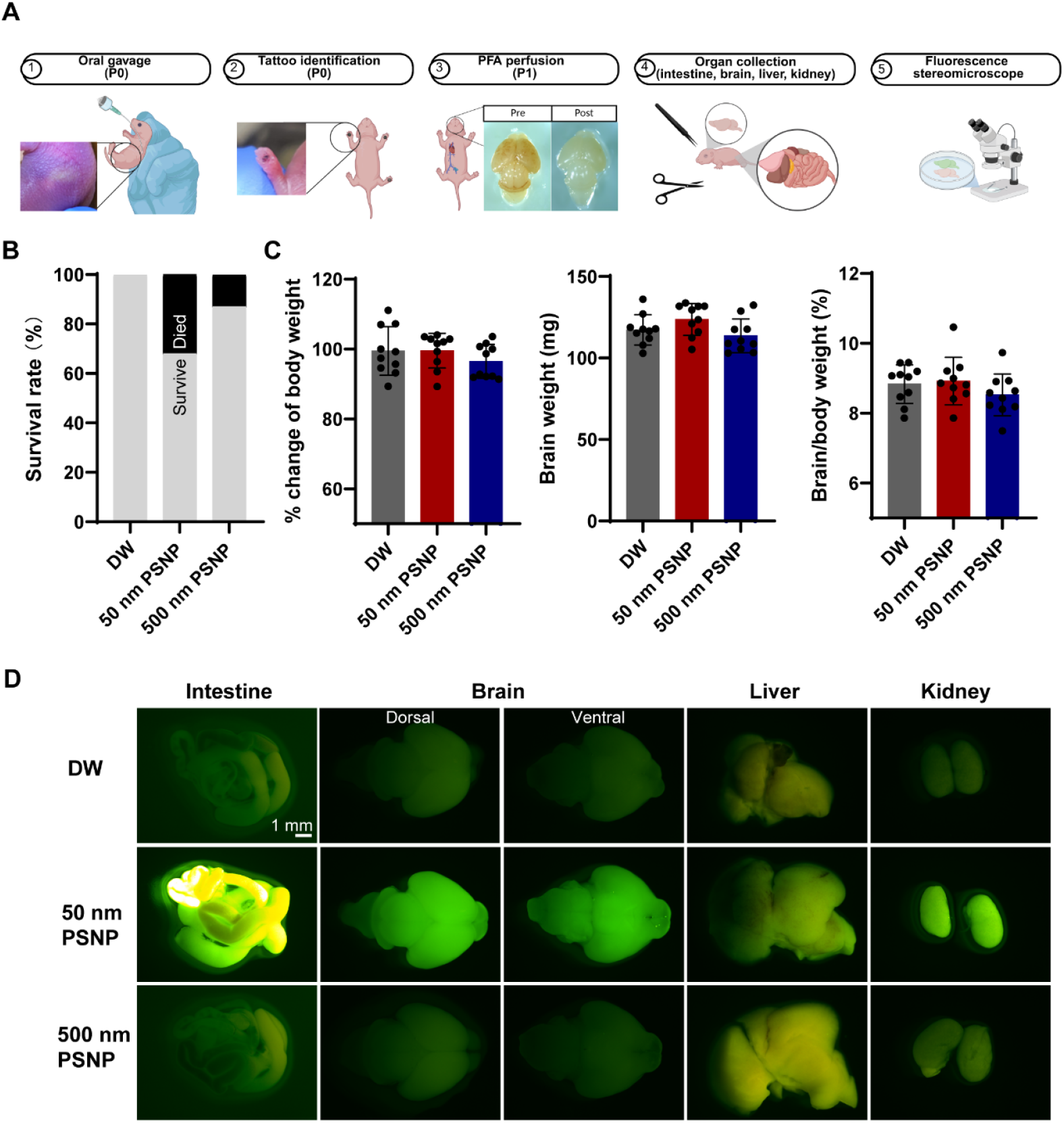
Size-dependent biodistribution of orally administered fluorescent polystyrene nanoplastics in neonatal mice. (A) Schematic of the experimental workflow: P0 mice received oral gavage with PSNPs or DW; organs were collected 24 h post-exposure for imaging (n = 5 per sex per group, derived from 6–8 dams). Created in BioRender (Smith, J., 2025; BioRender.com/c248457). (B) Survival rate per treatment group; neonatal mice were randomly assigned from 15 dams. Group sizes: DW (n = 18), 50 nm PSNP (n = 25), 500 nm PSNP (n = 16). (C) Body weight change (%) from P0 to P1, P1 brain weight, and P1 brain-to-body weight ratio in mice that survived exposure (n = 10 per group, derived from 6–8 dams). No significant differences were detected in body weight, brain weight, or brain-to-body weight ratio (one-way ANOVA, p > 0.05 for all comparisons). (D) Fluorescence stereoscope images of neonatal mouse organs showing NP accumulation in the intestine, brain, liver, and kidney. Compared with 500 nm PSNPs and DW controls, 50 nm PSNPs exhibited greater retention, particularly in the intestine. Scale bar = 1 mm. PSNP, polystyrene nanoplastic; P0, postnatal day 0; DW, distilled water; ANOVA, analysis of variance.

At 24 h post-treatment, the dissected organs were imaged using a fluorescence stereoscope (see Methods for details), and brain weights were measured to evaluate potential neurodevelopmental effects. No sex-specific differences were observed in weight quantification or imaging, and mixed-sex samples were used for subsequent analyses (n = 5 per sex, per group). No significant differences in body weight change (%) from P0 to P1 [F (2, 27) = 0.9385, p = 0.4036], P1 brain weight [F (2, 27) = 2.705, p = 0.085], or P1 brain-to-body weight ratio [F (2, 27) = 1.150, p = 0.3316] were observed across the three groups (Figure 1C). Organs selected for region-specific imaging - based on their roles in nanoparticle uptake, processing, and barrier sensitivity - showed pronounced accumulation of 50 nm PSNP, including in the intestine, brain, liver, and kidney, whereas 500 nm PSNP exhibited markedly weaker signals across these organs (Figure 1D). Notably, fluorescence intensity appeared stronger in the brain than in the liver and was broadly comparable to levels detected in the kidney - organs commonly associated with nanomaterial accumulation (Wang et al., 2016; Xu et al., 2024; Takáč et al., 2025) - suggesting that small NPs may more easily cross physiological barriers and reach the brain during early development. This pattern was especially consistent in intestinal samples, which may reflect the slower clearance of smaller NPs during the early postnatal period. These size-dependent biodistribution patterns are broadly consistent with previous reports in adult rodent models, where smaller particles demonstrated greater translocation across physiological barriers, including the BBB and intestinal epithelium (Zhang et al., 2023). Notably, these findings resonate with recent human data showing the presence of MNPs in the liver, kidney, and particularly brain tissue (Nihart et al., 2025), although direct quantification was not performed in the current study. This alignment across developmental stages and species suggests that NP diameter is a key determinant of systemic uptake and organ-level retention, particularly during early life windows when barrier functions may be immature or underdeveloped.

### 3.2 Regional brain distribution visualized by LSFM

Given the greater organ-level uptake of smaller particles, we selected 50 nm PSNP for whole-brain imaging to characterize regional biodistribution and evaluate the performance of our tissue optical clearing and LSFM workflow for NP detection. Neonatal mice were orally administered 30 μL of 50 nm PSNP (2.5 or 12.5 mg/mL) or DW as a control at P0. Brains collected at P1 (24 h post-administration) were optically cleared using SeeDB2G (Figure 2A), a simple immersion protocol that preserves fluorescence signals while enabling deep-tissue visualization without sectioning. Cleared samples were subsequently imaged using LSFM (see Methods for details), providing a 3D map of NP distribution in brain tissues. LSFM fluorescence images confirmed successful PSNP detection at both concentrations, with markedly stronger signals in the 12.5 mg/mL group than in the 2.5 mg/mL and control groups (Figure 2B). The advantages of LSFM were further illustrated by 3D reconstructions of brains treated with 12.5 mg/mL PSNP, visualized from ventral, lateral, and oblique angles, along with a heatmap visualization of fluorescence intensity, enabling high-resolution assessment of signal localization near ventricular boundaries (Figure 2C, Supplemental Video 1). This integrated approach demonstrates the potential of combining tissue clearing and LSFM to investigate complex biodistribution patterns without the need for serial sectioning.

**Figure 2.**
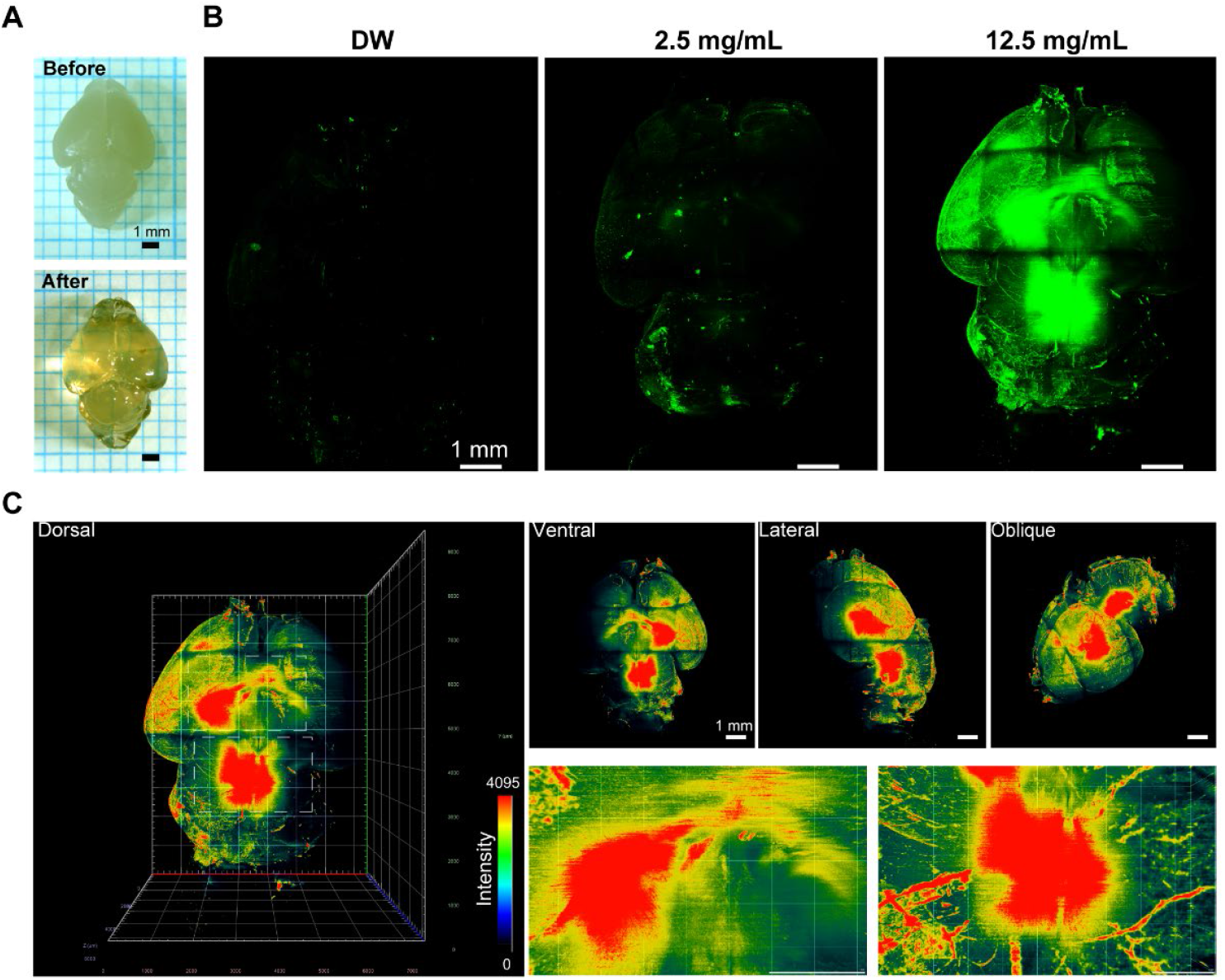
Regional distribution of fluorescent polystyrene nanoplastics in neonatal mouse brain visualized by LSFM. (A) Brightfield images of neonatal (P1) mouse brain before and after optical clearing using SeeDB2G, indicating successful clearing for deep-tissue imaging. (B) LSFM fluorescence images of cleared brains from mice orally treated at postnatal day 0 (P0) with 50 nm PSNPs (2.5 or 12.5 mg/mL) or DW as a control, imaged 24 h post-administration. (C) Three-dimensional LSFM reconstructions of cleared brains treated with 50 nm PSNPs (12.5 mg/mL), shown from ventral, lateral, and oblique views, with heatmap visualization of fluorescence intensity. Bottom panels show magnified regions highlighting strong signal localization near ventricular boundaries. Scale bar = 1 mm. LSFM, light-sheet fluorescence microscopy; PSNP, polystyrene nanoplastic; P1, postnatal day 1.

### 3.3 Two-dimensional section-based visualization and quantitative analysis using LSFM

Considering the anatomical complexity of the brain, LSFM enables analysis of both 3D reconstructions and two-dimensional sections extracted from the same dataset, enabling direct comparison with conventional brain atlases and improving anatomical precision in signal localization. Sagittal sections revealed prominent fluorescence in the thalamus and brainstem, with signal intensity increasing at higher concentrations (Figure 3A). Coronal sections further confirmed these patterns and highlighted additional signals in the cerebellum and cortex (Figure 3B).

**Figure 3.**
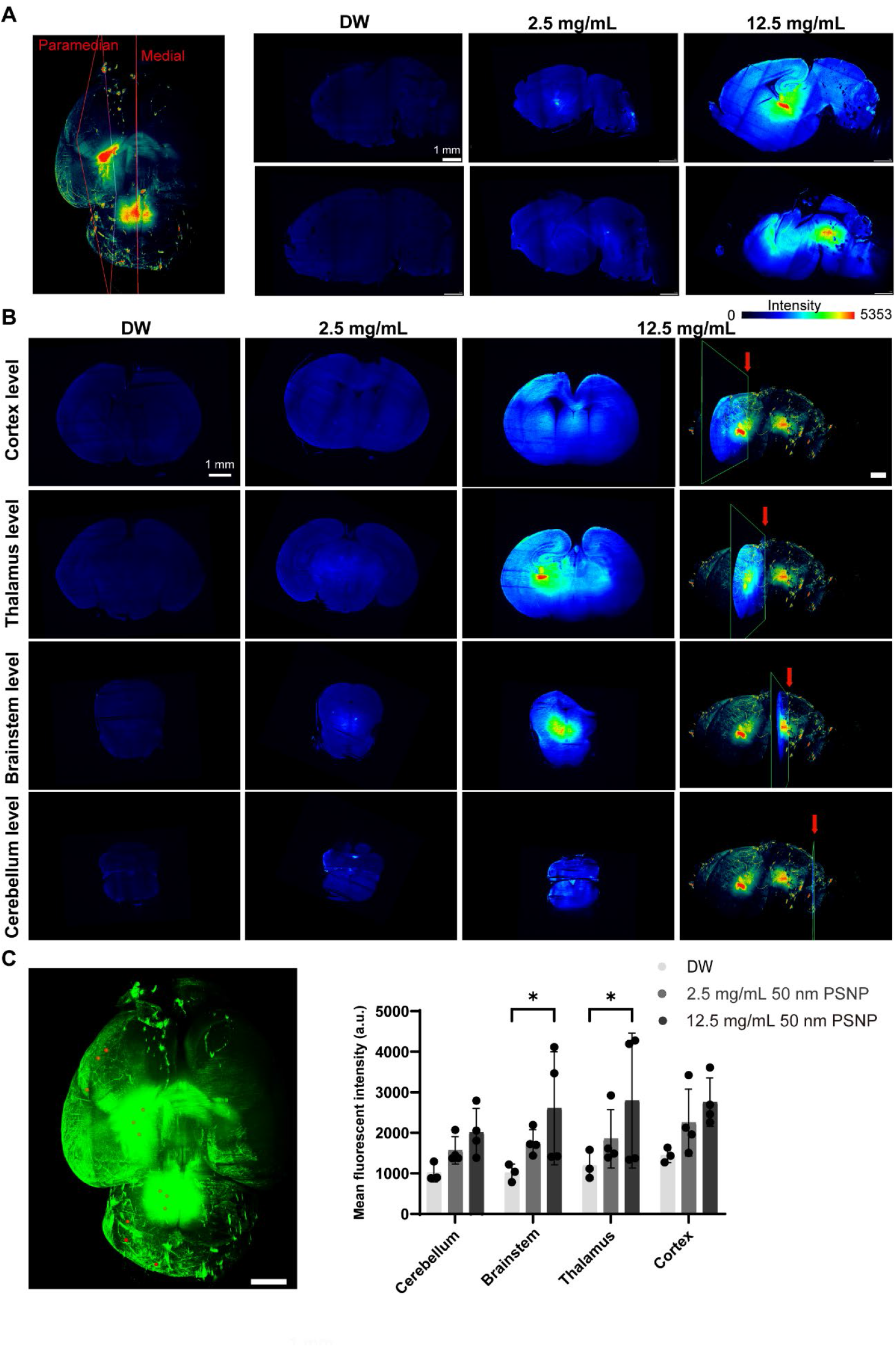
LSFM-based 2D sectional visualization and 3D quantification of PSNP in neonatal mouse brain. (A) Representative paramedian and medial sagittal LSFM sections of optically cleared neonatal mouse brains collected 24 h after oral administration at postnatal day 0 (P0) with 50 nm PSNP at 2.5 mg/mL or 12.5 mg/mL, or DW as a control. (B) Coronal sections extracted from LSFM datasets at cortex (section 073, 2.19 mm), thalamus (section 133, 3.99 mm), brainstem (section 189, 5.67 mm), and cerebellum (section 249, 7.47 mm) levels, illustrating concentration-dependent accumulation of PSNP. In the 12.5 mg/mL group, corresponding 3D reconstructions are provided, with arrows indicating the precise locations of each coronal section within the whole-brain view. Stronger signals are evident in the thalamus and brainstem at 12.5 mg/mL. (C) 3D LSFM rendering of a cleared brain treated with 12.5 mg/mL PSNP and quantitative analysis of mean fluorescence intensity within spherical ROIs (100 µm diameter) across four brain regions (cerebellum, brainstem, thalamus, cortex). Data are presented as means ± SDs; *p < 0.05 compared to DW (two-way ANOVA with Tukey’s post hoc test). Scale bar = 1 mm. LSFM, light-sheet fluorescence microscopy; PSNP, polystyrene nanoplastic; ANOVA, analysis of variance; SD, standard deviation.

Based on observable fluorescence patterns (Figure 3C), PSNP signals were broadly distributed throughout the brain, with the highest accumulations in the thalamus and brainstem at 12.5 mg/mL (two-way ANOVA: F(2, 32) = 9.891, p = 0.0005; Tukey’s post hoc: thalamus p = 0.0376, brainstem p = 0.0389). Similar— though less consistent—trends were observed at 2.5 mg/mL. Although increases in cortical and cerebellar signals were observed at both doses, the differences were not significant (adjusted p > 0.10). Notably, all regions exhibited dose-dependent increases in fluorescence, suggesting concentration-dependent accumulation despite interregional variability.

## 4. Discussion

This study introduces an integrated workflow combining SeeDB2G tissue optical clearing with LSFM to enable whole-brain, 3D visualization of NP biodistribution in intact neonatal mouse brains. Compared with conventional section-based imaging, LSFM allows volumetric mapping without slicing, preserving anatomical continuity and spatial context, which is crucial for interpreting regional particle localization in complex organs such as the brain. Using this workflow, we achieved deep-tissue imaging with sufficient fluorescence retention to support qualitative analyses.

Although the current workflow relies on fluorescently labeled NPs, limiting direct application to unlabeled environmental samples, it provides an essential foundation for 3D mapping of NP biodistribution. Future studies should incorporate environmentally relevant particles (Kochanek et al., 2025), which are far more heterogeneous than model PS beads in terms of polymer type, morphology, size, surface charge, and weathering state. In particular, polyethylene (PE) is the most frequently detected plastic pollutant in biological samples (Tiago et al., 2023). Addressing this complexity requires advanced label-free imaging strategies, such as atomic force microscopy, hyperspectral imaging (Nigamatzyanova et al., 2021), and optical photothermal infrared microspectroscopy (Macairan et al., 2025), which hold promise for in situ detection of unlabeled particles (Zhao et al., 2025). By combining LSFM volumetric mapping with these complementary approaches, researchers can construct a more complete and mechanistically informative atlas of nanoparticle uptake and clearance. Notably, the expanding availability of fluorescently labeled NPs (Merdy et al., 2024) of diverse polymers and sizes will enable more realistic comparative studies and strengthen hazard evaluations. Moreover, this workflow is broadly applicable to other organs (including the intestine, kidney, and liver) and species, supporting cross-tissue comparisons of nanoparticle translocations.

We observed size-dependent biodistribution following oral administration, with 50 nm PSNPs showing stronger organ-level signals than 500 nm PSNPs. Using the clearing and LSFM workflow, we further mapped widespread brain accumulation, with the strongest signals in the thalamus and brainstem, and clear dose-dependent increases across regions. Accumulation in these areas suggests potential entry via the choroid plexus-CSF pathway, followed by translocation across the ependymal CSF-brain barrier into the parenchyma, although contribution from the BBB cannot be excluded. These regions are adjacent to the fourth and third ventricles, where CSF is secreted by the choroid plexus. Given the developmental specialization of the blood-CSF barrier (Liddelow, 2015) - characterized by mature tight junctions and active transcellular transport - CSF-mediated entry is plausible. Once in the CSF, the relatively permissive ependymal layer may facilitate diffusion into surrounding brain tissue, particularly during the neonatal stages. Alternatively, the observed localization may reflect clearance (Madadi et al., 2024) via CSF circulation and glymphatic pathways (Liu et al., 2022), particularly for nanoparticles <100 nm (Gu et al., 2020). The potential interplay between entry and clearance mechanisms highlights an important avenue for future research.

Beyond tissue-level mapping, cell-type specificity remains a crucial next step. Prior evidence indicates that microglia exhibit the highest capacity for internalizing small PSNPs, followed by neural stem cells, whereas differentiated neurons and astrocytes show comparatively low uptake (Ito et al., 2025). indicating that particle size and cell identity strongly influence internalization, and regional accumulation may reflect the developmental cell-type composition. Consistent with our tissue-level findings of strong thalamic signals, Allen in situ atlas data indicated that progenitor markers such as Sox2 are expressed in this region during our study window, potentially explaining the regional differences in NP uptake. Extending the workflow to include cell-type markers will help identify the cellular contributors to regional nanoparticle accumulation. Collectively, these results underscore the value of combining optical clearing (SeeDB2G) with LSFM for the whole-brain NP biodistribution in neonatal mice. The workflow delivers anatomical continuity, volumetric context, and sensitivity to regional differences while acknowledging current limitations (fluorescent labeling and model particle choice). Using this approach, we showed that orally administered NPs cross neonatal physiological barriers and accumulate in the brain in a size-dependent and region-specific manner, with the strongest signals in the thalamus and brainstem.

## 5. Author Contributions: CRediT

**Yang Mi**: Writing – original draft, Writing – review & editing, Methodology, Visualization, Validation, Formal analysis, Data curation, Conceptualization, and Investigation. **Tomohiro Ito**: Writing – review & editing, Investigation, and Methodology. **Kosuke Tanaka**: Writing – review & editing, Investigation, Methodology, and Funding acquisition. **Osamu Udagawa**: Writing – review & editing, and Funding acquisition. **Masaki Kakeyama**: Writing – review & editing, and Funding acquisition. **Yasuo Tsutsumi**: Writing – review & editing, and Funding acquisition. **Fumihiko Maekawa**: Writing – review & editing, Conceptualization, Methodology, Resources, Funding acquisition, Supervision, and Investigation.

## 6. Acknowledgments

The authors thank M. Fujii and M. Matsumoto (National Institute for Environmental Studies) for their secretarial assistance. We would like to thank Editage (www.editage.com) for the English language editing.

## 9. Data availability

Data will be made available on request.

## 10. Declaration of generative AI and AI-assisted technologies in the writing process

During the preparation of this work the authors used Copilot in order to improve language refinement and readability. After using this tool, the authors reviewed and edited the content as needed and take full responsibility for the content of the published article.

