## Supplementary material for "Whole-Tissue Distribution Analysis for Visualization of Nanoplastics in the Mouse Brain": Supplimental Video 1: Supplemental Video_caption251223.pdf

**Supplemental Video 1 caption**

**Supplemental Video 1. Three-dimensional LSFM reconstruction of an optically cleared neonatal mouse brain (P1) treated with 50 nm PSNPs (12.5 mg/mL).** The video shows the rotational views and zoomed-in visualizations of the fluorescence signals.
